# Treadmill exercise promotes retinal astrocyte plasticity and protects against retinal degeneration

**DOI:** 10.1101/2021.06.07.447392

**Authors:** Katie L. Bales, Alicia S. Chacko, John M. Nickerson, Machelle T. Pardue, Jeffrey H. Boatright

## Abstract

Exercise has been shown to be an effective neuroprotective intervention that preserves retinal function and structure in several animal models of retinal degeneration. However, retinal cell morphology and cell types governing exercise-induced retinal neuroprotection remain elusive. Previously, we found that the protective effects of exercise in animal models of retinal disease were accompanied by increased levels of circulating and retinal brain derived neurotrophic factor (BDNF) and required intact signal transduction with its high-affinity receptor, tropomyosin kinase B (TrkB). Studies of neurodegenerative diseases in the brain demonstrate that neurons and astrocytes express BDNF and TrkB. Additionally, astrocytes have been shown to alter their morphology in response to exercise. Here, we have investigated the role of retinal astrocytes as mediators of exercise-induced retinal neuroprotection in a light-induced retinal degeneration mouse model (LIRD). We found that treadmill exercise in both our dim (control maintenance light levels) and LIRD groups promote increased retinal astrocytic population, GFAP expression, branching and endpoints, dendritic complexity, and promotes BDNF-astrocyte interaction. In contrast, LIRD animals that were inactive had significant reductions in all measured parameters. Our findings indicate that exercise is sufficient to rescue retinal astrocyte morphology in a LIRD model maintaining branching and dendritic arborization similar to retinal astrocytes that are not undergoing degeneration. These studies provide essential information to current knowledge gaps in regards to exercise-induced neuroprotection and will additionally provide knowledge in exercise intervention optimization as a rehabilitative method.

**Significance statement:** This study represents an essential step in determining the cell-types governing and morphological alterations elicited from exercise which may provide neural repair and protection. Similar to astrocytes in the brain, retinal astrocytes alter their morphology in response to exercise. Our studies demonstrate exercise promotes increased interactions between retinal astrocytes and neural growth factors in healthy retinas as well as in retinas undergoing degeneration, which may ultimately protect dying retinal neurons. These studies provide insight into the potential neuroprotective role astrocytes play in neurodegenerative diseases.

## Introduction

The rehabilitative and protective effects of aerobic exercise for prevention or treatment of neurodegenerative diseases have been well-documented in human and animal studies^(1-4)^. Further, clinical trials and retrospective studies suggest that exercise may have preventative and rehabilitative effects on visual outcomes in patients with retinal dystrophies, such as age-related macular degeneration and retinitis pigmentosa ^(4-7)^. There are minimal effective treatment options for patients with vision loss. These treatment options are often administered only after the disease has significantly progressed, and can be expensive and invasive^(8, 9)^. Exercise as a therapy for retinal degenerative disease offers a non-invasive, low-cost intervention that can be performed in any setting. In order to develop the most effective methods for exercise-mediated interventions to slow vision loss, it is imperative to identify the molecular mechanisms and cell-types modulating the protective effects of exercise. To date, the underlying processes and cell-types governing the neuroprotective benefits of exercise have yet to be fully characterized.

Previously, we found that exercise is neuroprotective in several animal models of retinal disease, displaying significantly reduced photoreceptor cell death and preserved retinal structure and function^(10-14)^. The protective effects of exercise were accompanied by increased levels of systemic and retinal brain derived neurotrophic factor (BDNF) and required signal transduction via the cognate receptor of BDNF, TrkB (tropomyosin receptor kinase B) ^(10, 14)^. Studies using rodent models of neurodegeneration demonstrate that BDNF and TrkB are expressed in both neurons and astrocytes and play significant roles in astrocyte morphological plasticity^(15-17)^. Additionally, it has been shown that exercise induces remodeling of astrocyte morphology in healthy brain tissue in animal models^(18-20)^.

As the most abundant cell type in the central nervous system, astrocytes are the glial cells responsible for a variety of tasks, including maintaining homeostasis, axon guidance, and synaptic support to control the blood brain barrier and blood flow^(21, 22)^. In the eye, astrocytes are essential in the development and function of the neural retina^(23)^. Retinal astrocytic bodies localize to the nerve fiber layer where they govern the microenvironment, provide mechanical support for damaged neurons, mediate neurotrophic function and contribute to the formation and maintenance of the blood-retinal barrier^(24)^. Astrocytes can exhibit extraordinarily complex morphology^(25)^. Although it is known that retinal glia experience morphological and functional changes with age and can be altered due to changes in the expression of trophic factors, few studies have focused on the pathways orchestrating how this complexity is maintained or altered in diseased states, especially within retinal degenerations and disease^(23)^. A growing body of literature suggests that BDNF and its high affinity receptor, TrkB, are expressed in neurons and astrocytes in the brain^(15, 17, 26)^. Additionally, other studies have shown that BDNF signaling is vital for proper astrocyte morphology in both development and adulthood^(15, 17, 26)^. Exercise offers a non-invasive and inexpensive method to increase BDNF signaling. Therefore, we hypothesize that astrocytes are a retinal cell-type mediating exercise-induced retinal neuroprotection.

In this study we investigated the potential role of retinal astrocytes as mediators of exercise-induced retinal neuroprotection through BDNF signaling mechanisms. We used a forced exercise paradigm of treadmill running as well as a light-induced model of retinal degeneration (LIRD) in order to control the amount of exercise as well as the onset of vision loss, respectively. We found in animals that were undergoing retinal degeneration that did not exercise (inactive+LIRD), retinal astrocytes had a significant decrease in branching, dendritic endpoints, dendritic arborization. Meanwhile, in animals that received exercise and were undergoing retinal degeneration (active+LIRD), retinal astrocytes had morphological similarities (branching, dendritic endpoints and dendritic arborization) to animals with healthy retinas (inactive+dim and active+dim). Through a proximity ligase assay (PLA), an immunodetection method used to identify two proteins that are close in space with high sensitivity and specificity, we found that exercise promotes extracellular BDNF-astrocytic interaction in active groups (both dim and LIRD) compared to inactive groups. These data support the potential role of retinal astrocytes as mediators of exercise-induced retinal neuroprotection through BDNF signaling mechanisms.

## Methods and Materials

### Animals

All experimental procedures were approved by the Institutional Animal Care and Use Committee of the Atlanta Veterans Affairs Healthcare System and conducted in accordance with the Association for Research in Vision and Ophthalmology Statement for the Use of Animals in Ophthalmic and Vision Research. Adult BALB/c male mice were purchased from Charles River (8-10 weeks old; Wilmington, MA, USA) and housed under a 12:12 light:dark cycle with ad libitum access to standard mouse chow and water.

### Experimental design

Mice were randomly assigned to one of the following four groups: inactive+dim (n=24), active+dim (n=24), inactive+LIRD (n=24) and active+LIRD (n=24). Active groups ran on a rodent treadmill once daily at 10 meters per minute (m/min), 5 days per week for 2 weeks. Inactive groups were placed on a static treadmill for the same amount of time. On the day of LIRD, they exposed to toxic light within 30 minutes of the end of the treadmill session. Following 1 additional week of treadmill running, electroretinography (ERG) was performed to assess retinal function (**Figure 1**). Following 2 final days of exercise, mice were exercised in a staggered fashion so that each mouse could be euthanized immediately after the end of the 1 hour of treadmill session. Mice were euthanized via CO_2_ and secondary cervical dislocation, eyes were enucleated for retinal flat mounts.

**Figure 1.**
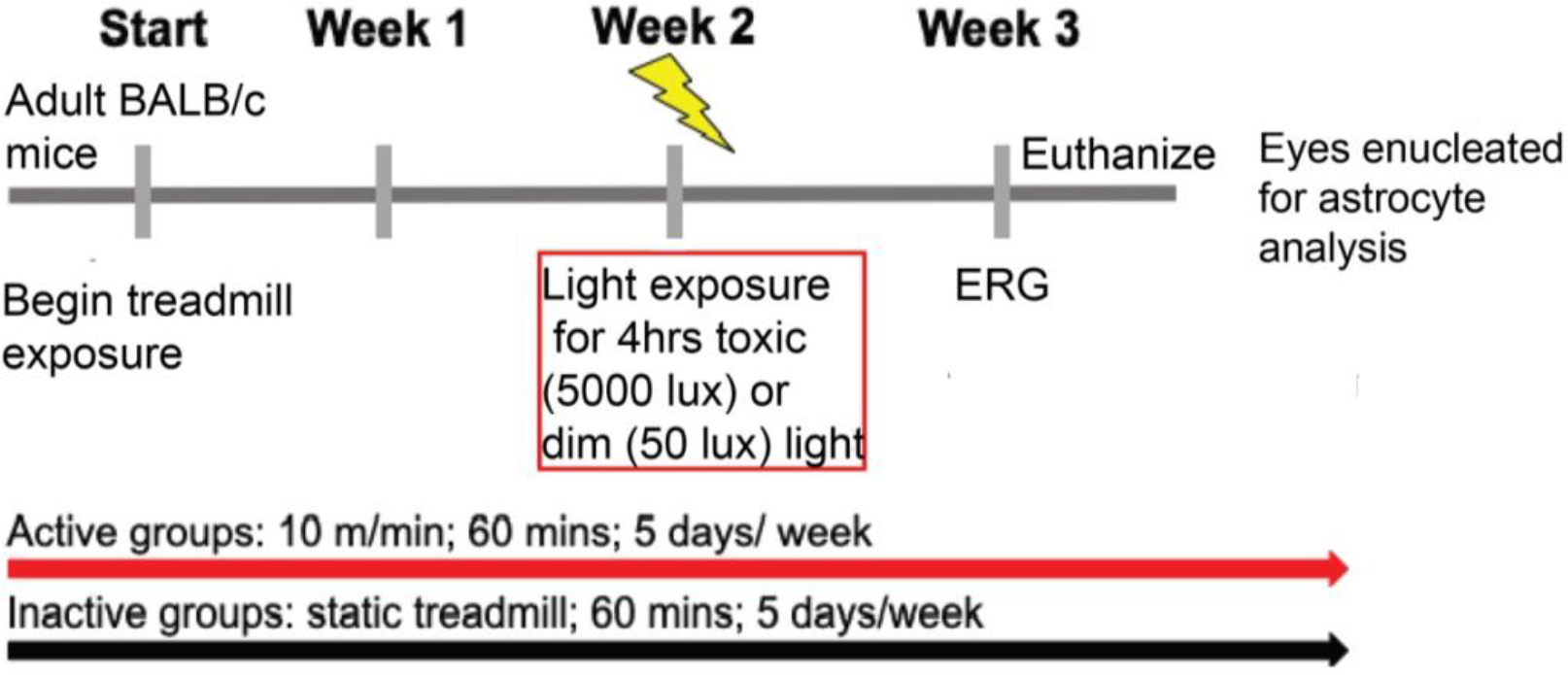
Experimental timeline for treadmill exercise and light-damage. Active groups were exercised by treadmill 1 hour a day for two weeks at a speed of 10m/min, meanwhile inactive groups were placed on static treadmills for the same duration. Light induced retinal degeneration (LIRD) occurred during the second week of exercise using light exposure of 5000 lux. Following retinal functional testing by electroretinography (ERG) in the third week mice were euthanized and retinal tissue was collected for histology and immunofluorescence.

### Exercise regimen and light exposure

In accordance with previous studies, active mice ran 60 minutes per day between 10 AM and 12 PM on treadmills equipped with electric shock grating (Exer-3/6; Columbus Instruments, Columbus, OH, USA) ^(10, 13)^. The grating inactivated if the mice received 10 shocks in a single session. Before daily treadmill sessions began the mice were trained for 5 to 10 minutes at a pace of 5 to 10 m/min for 2 consecutive days on the treadmill. The mice learned to maintain running within the first few days of treadmill exposure, receiving very few shocks after the first 1 to 2 days. Inactive mice were placed on static treadmills. Following 2 weeks of exercise, the mice were exposed to typical laboratory lighting (50 lux; dim) or toxic light (5,000 lux; LIRD) for 4 hours using a light emitting diode (LED) light panel (LED500A; Fancierstudio, Hayward, CA, USA). For light exposure animals were individually housed in shoebox containers with the LED light panel placed above as previously described^(13)^. Room and light box temperatures were closely monitored to ensure animal welfare.

### Electroretinography (ERG)

At 1 week following LIRD induction, retinal function was measured with a commercial ERG system (Bigshot; LKC Technologies, Gaithersburg, MD, USA) as previously described^(10, 13)^. After overnight dark adaptation, mice were anesthetized (ketamine [80 mg/kg]/xylazine [16 mg/kg]). The depth of anesthesia was considered appropriate to proceed when whisker movement had ceased and toe pinch reflex was absent. All procedures were performed under dim red light. The pupils were dilated (1% tropicamide; Alcon Laboratories) and corneas were anesthetized (1% tetracaine; Alcon Laboratories, Ft. Worth, TX, USA). Body temperature was maintained with a heating plate (ATC 1000; World Precision Instruments, Sarasota, FL, USA) for the duration of the session. The ERG protocol consisted of an eleven-step series of full-field flash stimuli produced by a Ganzfeld dome under both dark-adapted (−3.4 to 1.5 log cd s/m^2^) and light adapted conditions (2.0 log cd s/m^2^ 6.1 Hz flicker with a 30 cd/m^2^ background light). Custom gold-loop wire electrodes were placed on the center of each eye through a layer of 1% methylcellulose to measure the electrical response of the eye to each flash. Platinum needle reference electrodes (1cm; Natus Medical Incorporated, Pleasanton, CA, USA) were inserted subcutaneously in each cheek. Post-measurement, mice were given an IP injection of atipamezole (1 mg/kg) (Antisedan, Zoetis, Parsippany, NJ, USA) to counteract effects of xylazine^(27)^, saline eye drops and allowed to recover on a heating pad (37°C) before being returned to housing. ERG responses were measured for both eyes and averaged together.

### Retinal Flat Mount Immunofluorescence

Following treadmill exercise and our LIRD experimental design as described above, mice were euthanized and eyes enucleated for retinal flat mount preparation. Retinal flat mounts were permeabilized with 0.1% Triton X-100 in PBS. Retinal flat mounts were blocked in 5% normal donkey serum in PBS with 0.01% sodium azide and 0.3% Triton X-100 and primary incubations were in 5% normal donkey serum in PBS with 0.01% sodium azide. Retinal flat mounts were washed with PBS. Primary antibody incubations were performed for 24 hours at 4°C using an anti-glial fibrillary acidic protein (GFAP; Abcam, ab53554; 1:250 dilution) antibody. Secondary antibody incubations were performed for 1 hour at room temperature using Alexa Fluor 488-conjugated Donkey anti-Goat (ThermoFisher, A-11055; 1:500 dilution). Retinal flat mounts were mounted with ProlongGold (Cell Signaling, #8961). Representative regions on the superior-inferior axis and temporal-nasal axis each measuring 0.227mm^2^, were analyzed from each retina. This ensured representative sampling of the peripheral, medial and central zones of the upper, lower, temporal and nasal quadrants of each retina as previously described^(28)^. Retinal flat mounts were then imaged by confocal microscopy. Retinal tissue images were taken on an Olympus Fluoview1000 confocal microscope (Center Valley, PA) with a 60x objective and a Lumenera INFINITY 1-3C USB 2.0 Color Microscope camera (Spectra Services, Ontario, NY). All images were compiled and quantified using ImageJ software.

### Retinal astrocyte morphology analysis

Retinal flat mounts were fixed and labeled with the astrocytic marker GFAP as described above. Prior to image acquisition the experimenter was blinded to experimental conditions and animal group IDs. Confocal images were acquired as described above. Astrocyte branching and dendritic endpoint quantifications were assessed as described utilizing the AnalyzeSkeleton software plugins from FIJI software^(29)^. Sholl analysis was used to quantify the amount and distribution of astrocytic arborization at increasing radial distances through Sholl analysis plugin FIJI software^(30)^. In short, a line was drawn from the soma out to the furthest cellular process for each individual astrocyte. Sholl radii were set at starting 5μm from the start line with 1 μm increasing increments. Following analysis data were collected and binned at 5 μm intervals in GraphPad Prism 9.0.0 (San Diego, CA).

### Proximity Ligase Assay (PLA)

For *in situ* detection of BDNF and astrocyte interaction, proximity ligation assay (PLA; Duolink, Millipore Sigma) was performed in fixed retinal flat mounts as described above following the manufacturer’s instructions. Retinal flat mounts were permeabilized with 0.1% Triton X-100 in PBS. Retinal flat mounts were blocked in 5% normal donkey serum in PBS with 0.01% sodium azide and 0.3% Triton X-100 and primary incubations were in 5% normal donkey serum in PBS with 0.01% sodium azide. Retinal flat mounts were washed with PBS. Primary antibody incubations were performed for 24 hours at 4°C. Primary antibodies used for PLA detection were brain derived neurotrophic factor (BDNF; Novus Biologicals, 25928.11; 1:100 dilution) and glutamate transporter-1(GLT-1; Alomone, AGC-022; 1:100 dilution). Co-labeling for astrocytes was performed using the same anti-glial fibrillary acidic protein antibody described above (GFAP; Abcam, ab53554; 1:250 dilution). Secondary antibody incubation labeling GFAP positive cells was performed for 1 hour at room temperature using Alexa Fluor 488-conjugated Donkey anti-Goat (ThermoFisher, A-11055; 1:500 dilution) during the PLA probe incubation.

### Masking and Statistical analysis

Sample size was determined based on our previously reported data^(10, 11, 13)^. The personnel conducting assessments in experiments that required judgement were masked to the specific treatment group from which sampling arose. This included semi-automated marking of ERG a- and b-waves, image acquisition, semi-automated counting of GFAP positive cells in retina flat mounts, retinal astrocytic morphological quantifications and fluorescence quantification. Two-way ANOVAs with Tukey’s multiple comparison tests were performed using Graphpad Prism 9.0.0 (San Diego, CA). Data are displayed as mean ± SEM and results were considered significant if p<0.05.*p<0.05, **p<0.001, ***p<0.0001, ****p<0.00001.

## Results

### Treadmill running preserves retinal function in mice undergoing light-induced retinal degeneration

ERGs were performed on all experimental groups to assess retinal function (**Figure 2A-E**). Similar to our previous work, inactive+LIRD mice undergoing retinal degeneration had diminished ERG amplitudes compared to active+LIRD mice, which had statistically significant retinal function preservation as shown through a-wave (inactive+dim: 138.8μv ±36.44; active+dim: 156.2μv ±39.49; inactive+LIRD: 31.69μv ±6.17; active+LIRD: 73.68μv ±16.02; two-way ANOVA, F(9,60)= 5.435, p<0.0001), scotopic b-wave amplitudes (inactive+dim: 272μv ±40.60; active+dim: 286.4μv ±39.53; inactive+LIRD: 82.23μv ±9.70; active+LIRD: 153.4μv±28.31, two-way ANOVA, F(15,96)=2.380, p=0.0058) and photopic flicker b-wave amplitudes (inactive+dim: 57.83μv ±1.66; active+dim: 56.44 μv ±1.66; inactive+LIRD: 16.27μv ±1.34; active+LIRD: 29.66μv ±1.09) two-way ANOVA, F(3,39)=188.4, p<0.0001).

**Figure 2.**
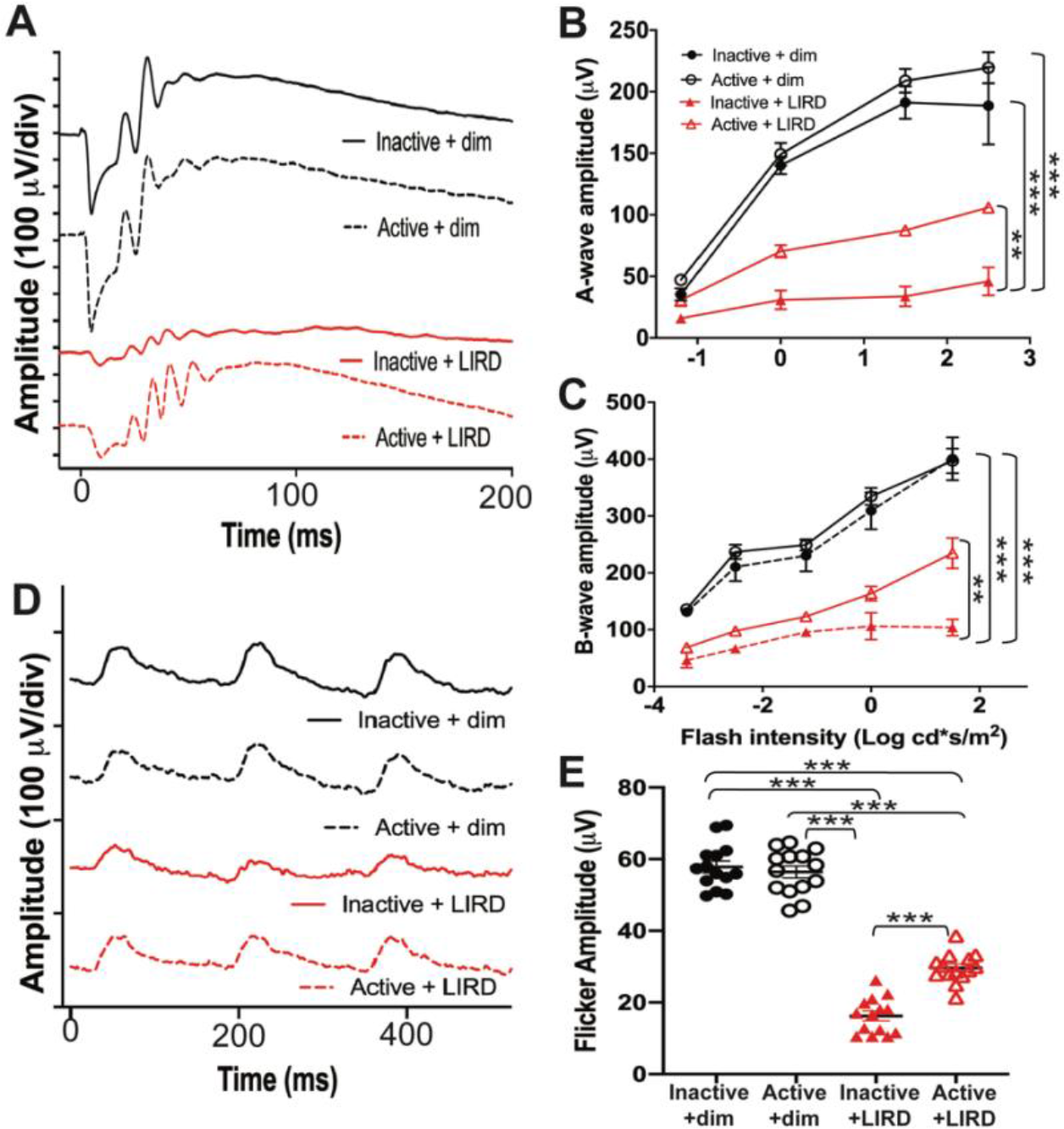
Treadmill exercise preserves retinal function in BALB/c LIRD mouse model and increases retinal BDNF expression. To ensure retinal neuroprotection was achieved, electroretinography (ERG) recordings were performed to measure retinal function non-invasively. Representative ERG waveforms are shown from maximum dark-adapted stimuli (1.5 log cd s/m^2^; **A**) and light-adapted flicker (2.0 log cd s/m^2^; **D**). ERGs revealed active+LIRD mice had significant preservation of a-wave (**B**), b-wave (**C**) amplitudes and flicker amplitudes (**E**) compared to inactive+LIRD mice. A- and b- waves show rod photoreceptor and inner retina function respectively, meanwhile flicker tests cone photoreceptor pathway function. **p<0.005; ***p<0.001. Values are mean ± SEM.

### Treadmill exercise alters retinal astrocyte density and morphology

We observed that individual retinal astrocytes from active groups had increased GFAP expression (inactive+dim:12.65±0.45; active+dim: 16.17±0.60; inactive+LIRD: 6.63±0.28; active+LIRD: 13.79±0.57; two-way ANOVA, F(3,57)=67.77, p<0.0001; **Figure 3E)** as well as increased retinal astrocyte populations throughout all four retinal quadrants (increased number of positive labeled astrocytes per image; inactive+dim: 6.12±0.21; active+dim: 8.8±0.25; inactive+LIRD: 3.96±0.15; active+LIRD: 7.76±0.20; two-way ANOVA, F(3,72)=102.6, p<0.0001 **Figure 3F**).

**Figure 3.**
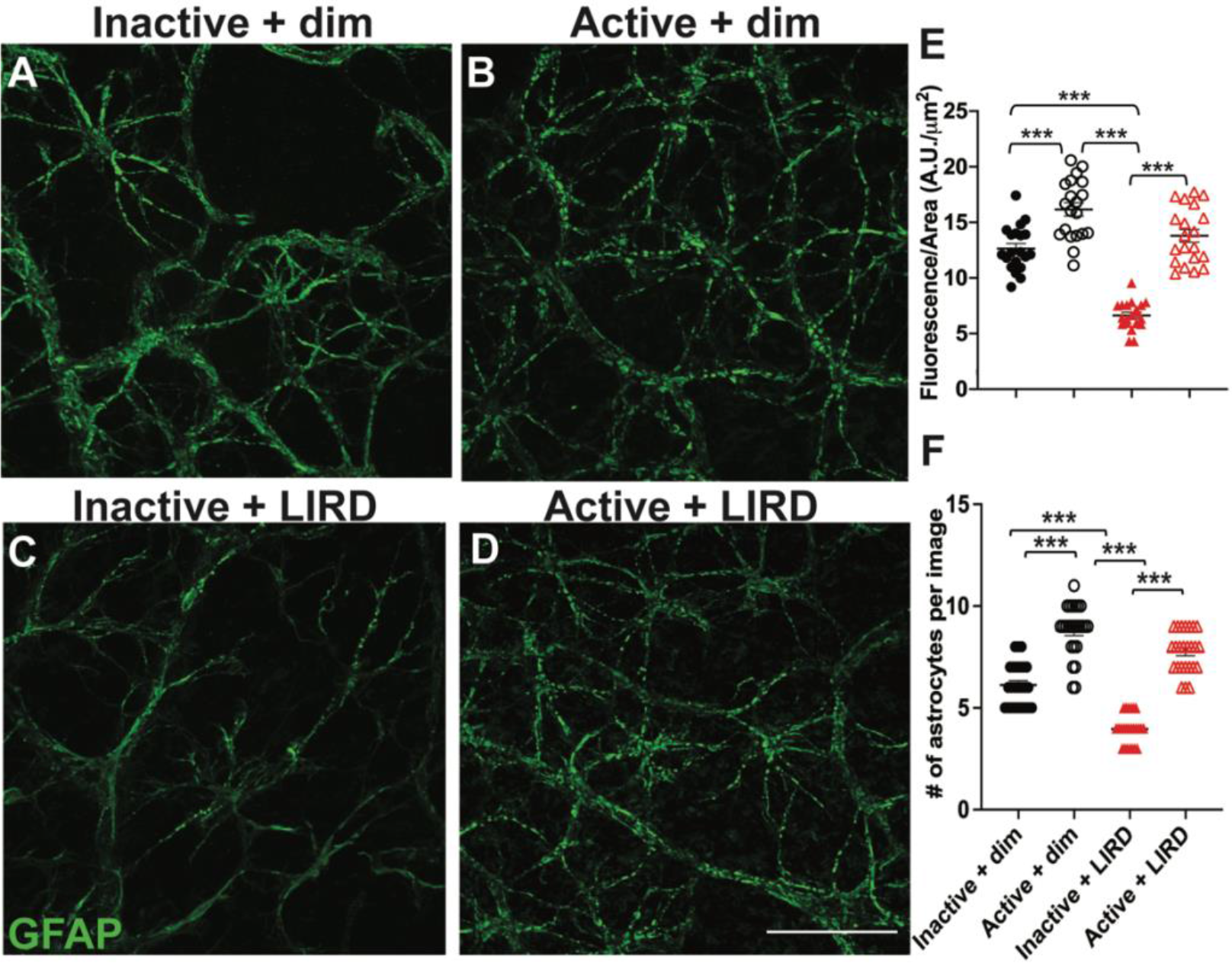
Retinal astrocyte density and expression is increased with exercise. Retinal flat mounts from inactive+dim (**A**), active+dim (**B**), inactive+LIRD (**C**), and active+LIRD (**D**) mice were stained for glial fibrillary acidic protein (GFAP, green). Cells positively labeled for GFAP were quantified by fluorescence (**E**) and number of astrocytes per image (**F**). Retinal astrocytes in both active groups had increased GFAP fluorescence as well as an increased in the number of positively labeled astrocytes present. A.U., arbitrary units, ****p<0.0001, scale bar=20μm. Values are mean ± SEM.

Astrocyte morphology was quantified using these GFAP-labeled flat mounts through the FIJI AnalyzeSkeleton Plugin (**Figure 4A-F**)^(29)^, revealing that active+dim and active+LIRD groups had increased astrocytic branching compared to respective inactive groups (inactive+dim:93.35±6.45; active+dim:155.1±6.40; inactive±4.40; active+LIRD:93.4±3.47; two-way ANOVA, F(3,57)=81.88, p<0.0001) and had an increased number of dendritic endings (endpoint voxels) compared to respective inactive groups (inactive+dim: 40±1.84; active+dim:83±4.58; inactive+LIRD:9.11±1.11; active+LIRD: 47.72±2.04; two-way ANOVA, F(3,51)=118.7, p<0.0001).

**Figure 4.**
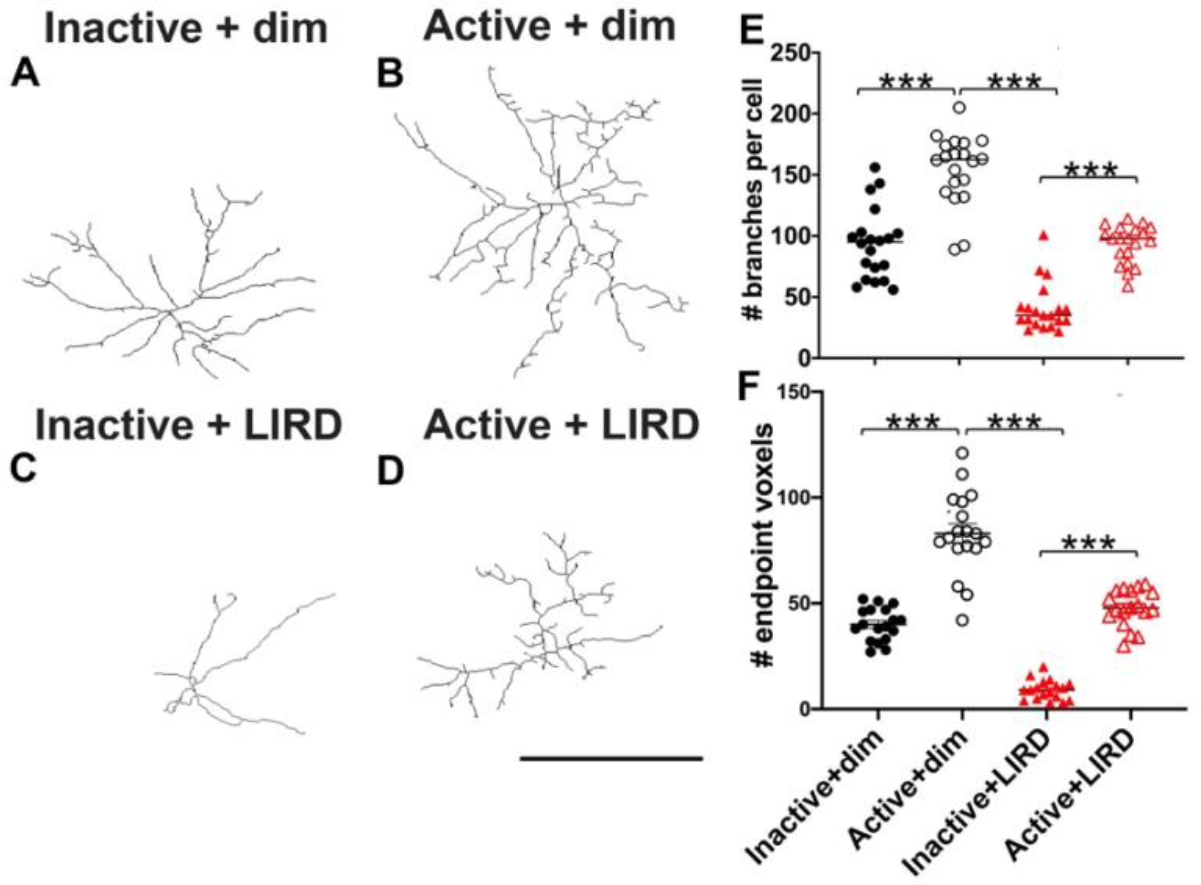
Active+LIRD have increased astrocytic branching and endpoint voxels. Retinal astrocytes from each experimental group were analyzed using Skeletonize plugin from FIJI (**A-D**). Astrocyte branches and endpoint voxels (dendritic endings) were then quantified (**E**,**F**) and two-way ANOVA multiple comparison analysis was performed. Active+LIRD retinal astrocytes appeared to be morphologically similar to inactive+dim astrocytes, in regards to branching and endpoint number and had no statistically significant differences (not significant; n.s.), whereas inactive+LIRD retinal astrocytes had a significant decrease in branching and endpoint voxels.****p<0.0001, scale bar= 20μm. Values are mean ± SEM.

### Retinal astrocytic dendritic complexity increased with treadmill exercise

We used Sholl analysis to quantify the effects of exercise on retinal astrocyte dendritic complexity. Sholl analysis revealed that inactive+LIRD retinal astrocytes had less dendritic arborization closer to the soma compared to inactive+dim, active+dim and active+LIRD groups as assessed by positive GFAP staining (**Figure 5A-E**; inactive+dim: 7.89±0.50; active+dim: 9.11±0.91; inactive+LIRD: 4.27±0.45; active+LIRD: 6.17±0.98; two-way ANOVA, F(3,479)=63.43, p<0.0001). Additionally, active+dim retinal astrocytes had a significant increase of dendritic arborization closest to the soma. Our data suggest that exercise increases retinal astrocytic branching, dendritic endpoints, and overall dendritic arborization, allowing increased signaling and synaptic connections in retinas undergoing degeneration.

**Figure 5.**
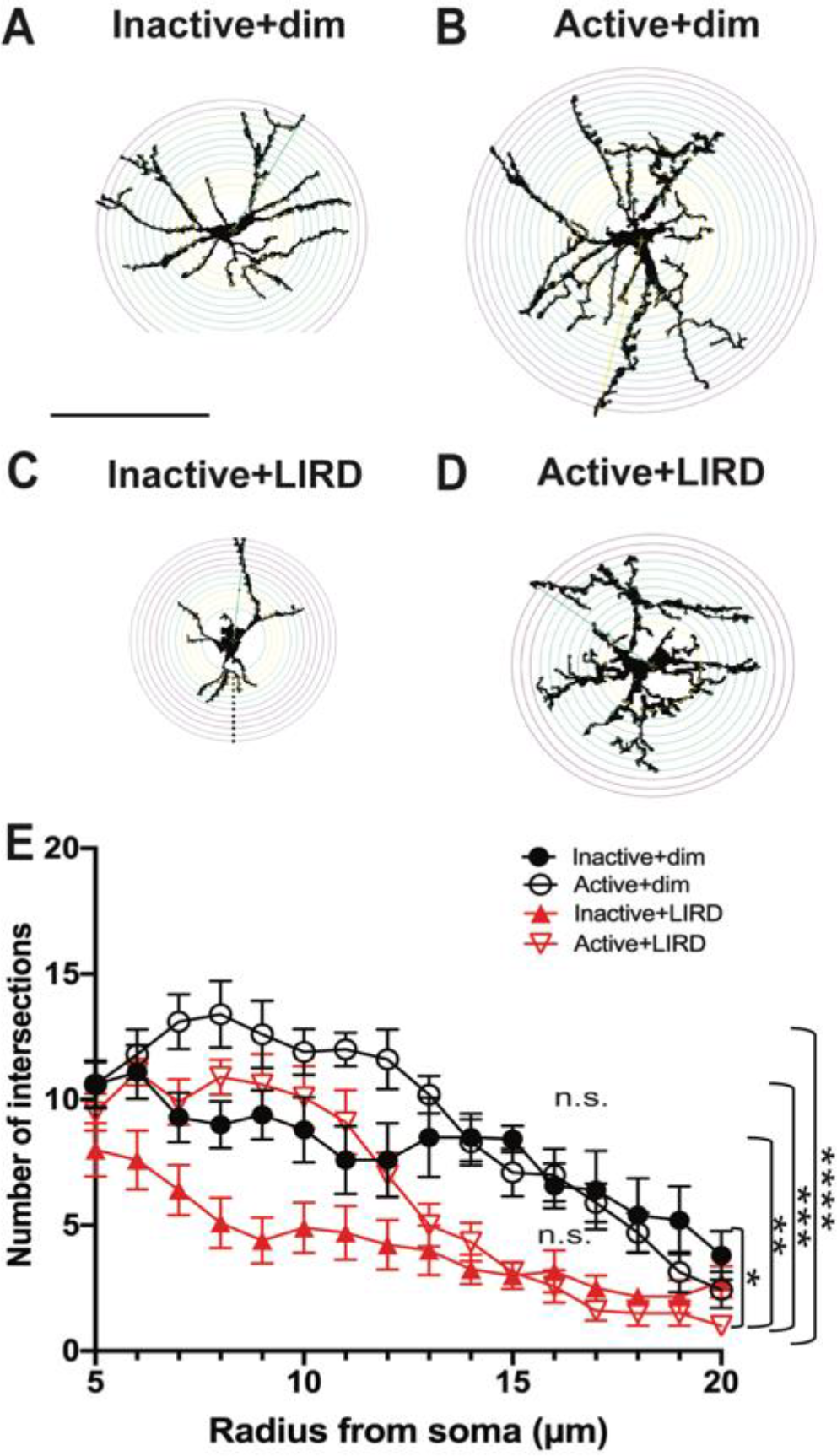
Sholl analysis reveals active+dim and active+LIRD astrocytes have increased complexity compared to inactive groups. Retinal astrocytes from each experimental group were analyzed using Sholl analysis plugin from FIJI (**A-D**). Astrocyte dendritic complexity was then quantified by counting the number of intersections from the soma (**E**). Two-way ANOVA multiple comparison analysis was performed. Retinal astrocytes from both active+dim and active+LIRD groups had significant increased dendritic complexity, with increased dendritic intersections occurring closer to the soma (7-10 microns from the soma) compared to inactive+dim and inactive+LIRD groups. N.s., no significance, *p<0.05, **p<0.01, ***p<0.005, ****p<0.0001, scale bar= 20μm. Values are mean ± SEM.

### Increased retinal BDNF-astrocyte interaction observed in treadmill exercised mice

To examine the interaction of BDNF and astrocytes *in situ*, we used the proximity ligation assay (PLA). PLA is an immunodetection method used to identify two proteins that are close in space with high sensitivity and specificity^(31, 32)^. Active groups (dim and LIRD; **Figure 6B and 6D**) showed a statistically significant increase in BDNF-GLT-1 interaction by PLA detection localized to astrocytes, whereas inactive groups (dim and LIRD, **Figure 6A and 6C**) displayed minimal PLA interaction occurring in retinal astrocytes (inactive+dim: 0.92±0.13; active+dim: 4.27±0.361; inactive+LIRD: 0.41±0.13; active+LIRD: 4.46±0.35; two-way ANOVA, F(3,57)=61.29, p<0.0001). BDNF-GLT-1 interaction appears to occur throughout the cell body and astrocytic branches.

**Figure 6.**
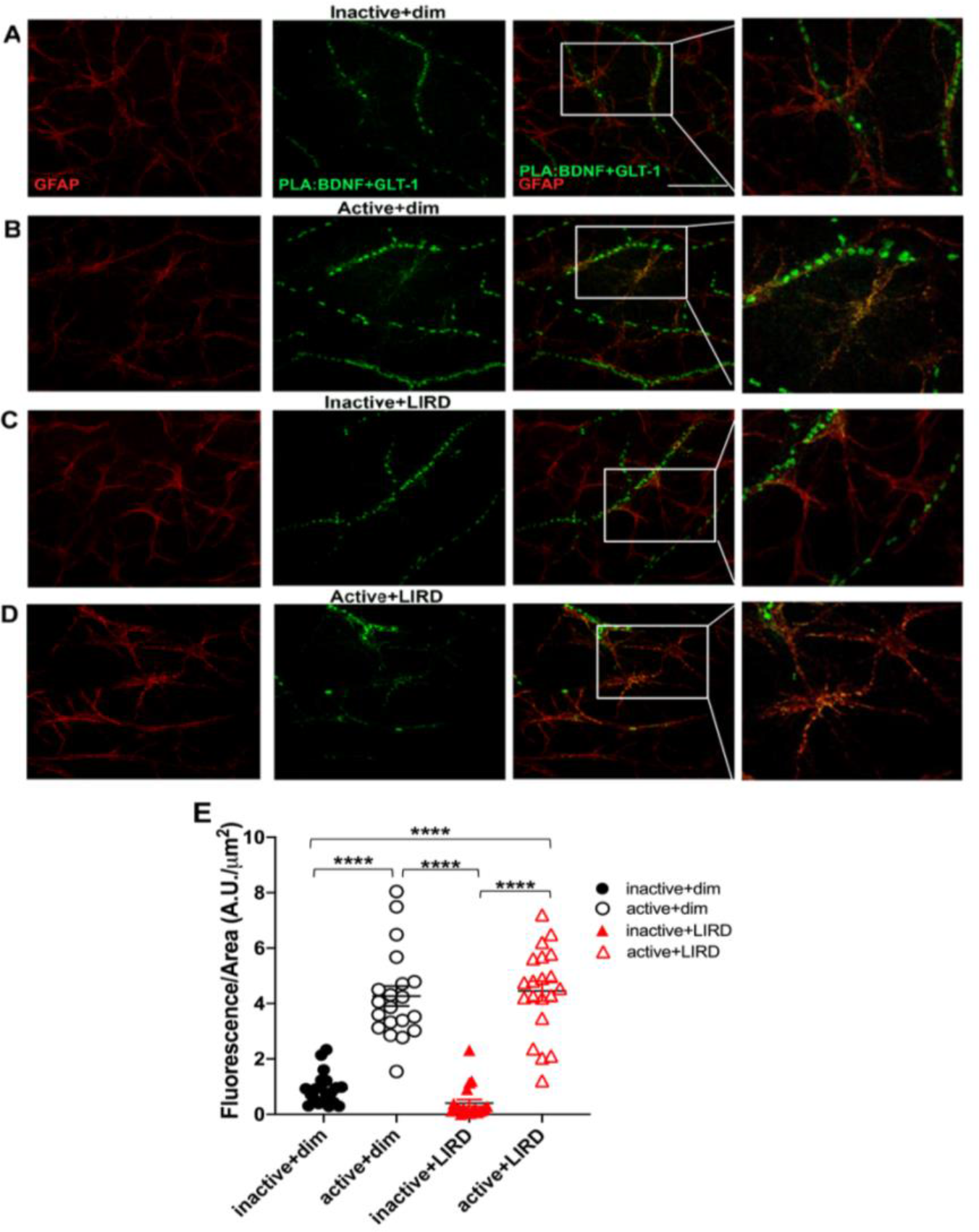
Treadmill exercise increases retinal astrocyte-BDNF interaction. Proximity Ligase Assay (PLA) was performed on retinal flat mounts from inactive (**A and C**) and active (**B and D**) dim and LIRD exposed mice using antibodies targeting BDNF and glutamate transporter-1 (GLT-1; green). GLT-1 is an extracellular membrane bound transporter expressed in astrocytes and endothelial cells. Co-labeling for astrocytes was also done using GFAP (red). Green fluorescence indicate BDNF-astrocyte interaction, yellow fluorescence signifies co-labeling of retinal astrocytes and BDNF. Quantification of PLA fluorescence reveal signification increased BDNF-GLT-1 (**E**). ****p<0.0001, scale bar=15 μm, white box signifies magnified region. Values are mean ± SEM.

## Discussion

In the retina, astrocytes have emerged as critical governors of retinal homeostasis and maintenance under both healthy and diseased conditions^(33)^. Retinal astrocytes respond to disease and injury, which leads to altered cell signaling and results in morphological and functional changes^(28)^. Our group has previously shown that exercise-induced retinal neuroprotection was associated with increased systemic and retinal BDNF expression^(10, 11, 14)^. These studies led us to investigate what retinal cell types might mediate BDNF signaling during exercise-induced retinal neuroprotection. Recently, brain studies in animal models have shown that astrocytes express both BDNF and its high-affinity receptor TrkB and change their morphology with BDNF expression^(17, 26)^. In this context, we investigated the potential role of retinal astrocytes in mediating BDNF expression in regards to exercise-induced retinal neuroprotection.

Using a forced exercise paradigm of treadmill exercise and a light-damage retinal degeneration model, we were able to control exercise duration and intensity, as well as the onset of retinal degeneration. Retinal astrocytes from active groups had increased GFAP expression as well as increased astrocyte population density compared to inactive groups. These data align with previous reports of exercise experiments using healthy brain tissue from animal models that show data indicating that astrocytes have increased GFAP expression and proliferation with exercise^(18-20)^. Morphological analyses revealed retinal astrocytes from inactive+LIRD mice had a significant decrease in astrocytic branching, dendritic endpoints and arborization. Meanwhile, active+LIRD mice had increased astrocytic branching, dendritic endpoints and arborization morphology statistically similar to inactive+dim mice.

These data show that treadmill exercise alters retinal astrocyte morphology not only in a healthy retina but also in a retina that is undergoing degeneration. From our statistical analysis, we were able to determine that active+LIRD retinal astrocytes were morphologically similar to inactive+dim mice. In order to get a more thorough understanding of how morphology effects function, future studies could evaluate astrocyte behavior to determine if active+LIRD retinal astrocytes also function similar to inactive+dim.

Performing a PLA assay, we were able to evaluate BDNF-astrocytic interaction, and found increased BDNF-astrocytic interaction in active retinal astrocytes compared to inactive groups^(31, 34)^. These data confirm that retinal astrocytes express BDNF and treadmill exercise promotes an increase in this interaction and or expression. Previously, it has been demonstrated that astrocytes express BDNF and its high-affinity receptor, tropomyosin kinase B (TrkB)^(17)^. Astrocytic BDNF and TrkB have also been shown to regulate severity and neuronal activity in the brain of neurodegeneration models^(15-17, 26)^. From this assay, we are also able to evaluate the cellular localization of BDNF-GLT-1 interaction in retinal astrocytes and found that is localizes to the cell body but also in astrocytic dendrites. The BDNF gene is processed at two different polyadenylation sites, which leads to mRNA transcription with two different length 3’ untranslated regions (UTRs; termed shorter 3’ UTR and longer 3’ UTR)^(35)^. These transcripts were found to differentially regulate neuronal dendritic arborization and are preferentially targeted to different parts of the cell (shorter 3’ to cell body and longer 3’ to cell body and dendrites)^(35)^. Therefore, treadmill exercise could be promoting BDNF expression within retinal astrocytes, stimulating increased cellular signaling, potentially providing retinal neuroprotection.

Future studies will evaluate which subclass of astrocytes are expressing BDNF (i.e. reactive or non-reactive) as well as other retinal cell types may be expressing BDNF as well as its receptor, TrkB, due to exercise and how systemic exercise potentially alters the cellular expression profiles of retinal cells. Our data indicate that retinal astrocytes may play an active role in exercise-induced retinal neuroprotection. We have not only shown that under retinal degeneration retinal astrocytes have diminished morphological complexity but additionally our data suggest that exercise can alter retinal astrocyte morphology, to the point where it is observationally similar to and quantitatively statistically indistinguishable from healthy retina. Additionally, we show that exercise increases BDNF-astrocytic interaction.

This study provides essential information for current knowledge gaps in regards to astrocyte morphology during retinal degeneration and their potential role in governing exercise-induced neuroprotection. These data provide insight to how whole-body exercise can influence retinal cell morphology, ultimately impacting cellular function and insight to specific cell types governing BDNF signaling mechanisms. Determining the retinal cell types governing molecular mechanisms underlying exercise-induced neuroprotection will provide vital information towards the development of the most effective methods for exercise-mediated interventions to slow vision loss in patients with retinal degenerations.

## Notes

### Competing Interest Statement

The authors have declared no competing interest.

